# New Finnlakevirus isolate FLiP-2 provides insight into the ecology of ssDNA phages in Flavobacterium hosts

**DOI:** 10.1101/2025.08.22.671703

**Authors:** Kati Mäkelä, Elina Laanto, Reetta Penttinen, Janne Ravantti, Kim Kreuze, Lotta-Riina Sundberg

## Abstract

Life cycle details or ecological impact are well characterized only for a few ssDNA phages. The *Finnlakeviridae* family includes one species, *Finnlakevirus FLiP*. Here, using the same *Flavobacterium* host and sampling location, we isolated a new strain designated FLiP-2, with 96,7 % genetic identity to the original isolate FLiP. To understand the ecology of this *Flavobacterium-*infecting phage species, we explored the host interactions of the two FLiP strains and a dsDNA Flavobacterium phage MaF61 under various conditions representing those encountered in their natural habitats in boreal lakes including different temperatures, anoxic conditions, and in the presence of different nutrients and antibiotics. While FLiP and FLiP-2 had similar virion stability outside the host, they exhibited significant differences in plaque morphology and infectivity. FLiP-2 could not replicate in the presence of ampicillin, whereas FLiP thrived even at high concentrations. Both strains of *Finnlakevirus FLiP* propagated better under or after stress exposure compared to MaF61. Additionally, *Finnlakevirus FLiP* plaques appeared far from the original infection site, particularly in response to stress, suggesting latent presence within a motile or filamentous bacterium. In conclusion, *Finnlakevirus FLiP* showed remarkable flexibility in host-interactions being well adapted to fluctuating conditions in boreal freshwaters.

## Introduction

Single-stranded DNA (ssDNA) phages exhibit remarkable genetic diversity and abundance in many types of environments worldwide, challenging the traditional view that tailed double-stranded DNA (dsDNA) phages dominate viral communities (Székely and Breitbart 2016; Tucker et al. 2011). Metagenomic studies have revealed the widespread distribution and prevalence of ssDNA phages, especially *Gokushovirinae*, which form a subfamily of *Microviridae* (Székely and Breitbart 2016; Hopkins et al. 2014). *Microviridae* genomes have been detected in samples originating from marine and freshwater ecosystems, mammalian including human gut, marine vertebrates and invertebrates, insects, reptiles, sediments and soils, among other sources (Kirchberger et al. 2022; Labonté et al. 2015). Phages belonging to the families *Microviridae* and *Inoviridae,* both of which were accepted to International Committee on Taxonomy of Viruses (ICTV) classification in 1978, have been studied more in depth compared to the more recently established ssDNA phage families *Plectroviridae* (ratified 2020), *Paulinoviridae* (2021) and *Finnlakeviridae* (2020). Phages belonging to *Microviridae* and *Finnlakeviridae* are icosahedral in morphology while the other three ssDNA phage families consist of tubular phages. *Finnlakeviridae* phages are *Flavobacterium*-infecting ssDNA phages with a lipid membrane inside a tailless capsid (Mäntynen et al. 2020; Laanto et al. 2017). In the *Finnlakeviridae* family, there is currently only one known species, *Finnlakevirus FLiP* (Mäntynen et al. 2020; Laanto et al. 2017), and the information on the life cycle details in this group is so far very limited (Mäkelä et al. 2024). In addition to the above-mentioned ssDNA families, there are unclassified ssDNA phage isolates such as phi48:2 and phi18:4 that infect *Cellulophaga baltica* (Holmfeldt et al., 2013) and *Flavobacterium* infecting phage phiCJT23, which shares remarkable structural similarity with *Finnlakevirus FLiP* (Kejzar et al. 2022). The presence of an internal lipid membrane has already been confirmed for phiCJT23 (Kejzar et al. 2022), and chloroform sensitivity of the virions also suggests lipids in phi18:4 (Howard-Varona et al. 2025).

Phages and their host bacteria are ubiquitous across diverse ecosystems, including extreme environments such as highly saline salt flats, acidic hot springs, and polar ice caps, highlighting their adaptability (Ramasamy et al. 2023; Wani et al. 2022; Schloter et al. 2000; Shu and Huang 2022; Fister et al. 2016). However, constant harshness of the environment causes unidirectional selective pressures for both phages and their hosts, while in some environments recurring and unpredictable fluctuations in abiotic and biotic conditions promote the evolution of diverse adaptive mechanisms like phenotypic plasticity, and the maintenance of genetic polymorphisms (Zhang et al. 2023). Even in comparably moderate conditions like boreal lakes, phages and their hosts encounter challenges related to substantial seasonal variations in environmental conditions. Protein capsids of phages protect their genome from environmental factors such as UV radiation, desiccation, temperature extremes, and enzymatic degradation, but selection for stress-resistant phenotypes still occurs (Gomez et al. 2022). However, impact of environmental stressors to phages often occurs through the altered metabolism and activated stress responses in the host (Zhang et al. 2023). In general, if the same phage species can be recovered from an ecosystem repeatedly even after a long period, it indicates that both the phage and at least one of its host bacteria have adapted well enough to the local conditions to survive, even if the conditions are fluctuating.

Environmental factors influence all the steps of the phage life cycle. Condition-dependent “decisions” begin immediately when a phage encounters a potential host. For example, variations in temperature and salt concentration may change the availability and conformation of phage receptors, affecting phage binding affinity to them as well as the rate of adsorption (Moldovan et al. 2007). After a successful genome entry, strictly lytic phages can only follow the lytic cycle, while in sub-optimal conditions temperate phages may adopt lysogenic cycles (Howard-Varona et al. 2017). A longer-term association with the host bacterium can be more beneficial under certain conditions, facilitating the transmission of phages into new environments via moving cells. Most importantly, under stressful conditions, phage-host interactions may become more mutualistic: bacterial antiviral systems may be downregulated (Feiner et al. 2015), and phage-mediated mechanisms to cope with stress are more active (Nanda et al. 2015; Argov et al. 2017). Furthermore, phage-mediated horizontal gene transfer and phage-encoded auxiliary metabolic genes may enhance bacterial stress resistance and biofilm formation (Obeng et al. 2016; Huang et al. 2021). Prophages are typically induced to enter the lytic cycle by environmental changes (Howard-Varona et al. 2017). The presence of antibiotics, reactive oxygen species, and extreme temperatures are examples of stressful conditions that may lead to the activation of the bacterial SOS response and a phenomenon known as stress-induced increase of phage production (Kim et al. 2018; Comeau et al. 2007). This process is commonly based on upregulated *recA* expression, which plays a crucial role in the SOS response by inhibiting key proteins in bacterial cell division, but stress conditions can lead to cell filamentation even without SOS response (Comeau et al. 2007; Kim et al. 2018). This results in an increased need for holins to reach concentrations sufficient to effectively work on the enlarged inner membranes of the larger-than-normal cells. Consequently, this leads to delayed lysis and increased time to produce phage particles per cell before lysis is possible (Kim et al. 2018; Kaur et al. 2012; S. B. Santos et al. 2009). Furthermore, environmental factors such as temperature, pH, and salt concentration may influence the timing and duration of the lytic cycle steps inside the cell (Fister et al. 2016; Jończyk et al. 2011).

In this study, we aimed to describe the influence of environmental factors on Finnlakevirus life cycle. We isolated a new *Finnlakeviridae* phage, which is a strain of *Finnlakevirus FLiP* (Laanto et al. 2017). Life cycle characteristics of the original isolate FLiP and the new isolate FLiP-2 were compared with a tailed dsDNA phage MaF61. To understand phage life cycles in their natural environments, we exposed strains of *Finnlakevirus FLiP* to various conditions that it might face in the boreal lake environment: large seasonal temperature fluctuations, changing nutrient compositions and concentrations, occasional depletion of oxygen and presence of antibiotics. To differentiate between direct impacts of these factors on phages and effects experienced through host responses, three *Flavobacterium* hosts were used. We found that despite the high sequence identity (96,7 %) between FLiP and FLiP-2, their life cycles were differently affected by the host strain and environmental factors. Infection patterns were different between these phages depending on nutrient composition and concentrations. FLiP was able to replicate efficiently even in high ampicillin concentrations, whereas FLiP-2 barely replicated even in lower concentrations. Most interestingly, *Finnlakevirus FLiP* showed a unique response to host stress - plaques spreading to the environment from the original infection site – suggesting Finnlakeviruses may be maintained in a latent form within stressed cells and recovered to infective cycle under improved conditions.

## Materials and methods

### Isolation of phages and bacteria

ssDNA *Finnlakevirus FLiP* has been isolated in September 2010 from lake Jyväsjärvi in Central Finland (Laanto et al. 2017). Host bacteria of FLiP (Mäkelä et al. 2024), *Flavobacterium* sp. strains B114, B167 (Laanto et al. 2011) and B330 (Laanto et al. 2017) were used to isolate new phages. Bacterial liquid cultures were grown with agitation (120 rpm, RT). All bacterial strains were stored in −80°C with 20 % glycerol.

Water samples of 5 liters from lake Jyväsjärvi (62°13’46.4"N 25°44’16.4"E) and water samples of 1 liter from other freshwaters in Finland (Supporting information Table S1) were collected to assess the presence of FLiP-like phages. Phage isolation was done only from lake Jyväsjärvi. A two-step prefiltering practice was applied to remove eukaryotes and particles like pollen and dust: sterilized, reusable fabric filters were first used for coarse filtering after which Whatman Chromatography paper (Grade 3 mm Chr, pore size ∼ 6-11 µm) was used to further filter out any larger particles. After prefiltering, water sample was filtered through 0.2 µm membrane filters (500 ml rapid-flow bottle top filter aPES membrane, 75 mm dia) to remove all cells. Viruses were concentrated from the filtrates using polyethylene glycol 6000 (10 % w/v) and NaCl (0.5 M) mixing for 30-90 mins at 6℃. Samples were centrifuged (Thermo Scientific Sorvall Lynx 6000 centrifuge, F9-6×1000 -rotor, 13900 x g, 60 min, 4°C) and the pellets were resuspended into 0.2 mM potassium phosphate buffer (pH 7.2). The concentrated phage sample (100 µl) was then plated with each of the bacterial strains using Shieh medium (Song et al. 1988) in a double layer plaque assay: 3 ml soft-agar [0.7 % (w/v)] was tempered into + 47°C, after which 100 µl of bacterial culture and 100 µl of phage sample were added and the mixture was poured onto an agar plate [1 % (w/v)]. Plates were incubated 1-2 days at RT, after which plaques were collected and each resuspended into 400 µl of Shieh medium. Newly isolated phages were subjected to three rounds of plaque purification using double layer plaque assay with the isolation host.

### Extraction of bacterial DNA from filter membranes

After filtering the freshwater samples, bacterial DNA was extracted from 0.2 µm filter membranes with DNeasy PowerLyzer PowerSoil Kit (Qiagen) according to instructions of the manufacturer. Diverging from the instructions, no soil samples were used but a half of the 0.2 µm filter membrane per extraction tube. Tissue Lyser II homogenizer (Qiagen) was used for detaching samples from the filter membranes and to disrupt the bacterial cells (frequency 25/s, 30 sec).

### PCR screening with FLiP primers

PCR (Supporting information) was used to screen if DNA extracted from the filters has similar sequences to FLiP major capsid protein (MCP) or replication initiation protein (Rep). Dream Taq polymerase was used according to instructions of the manufacturer. PCR products were run in an 1 % agarose gel.

### Phage lysate preparation

To perform the experiments, high titer (10^10^-10^11^ plaque forming units per milliliter, PFU ml^-1^) lysates of FLiP, FLiP-2 and MaF61 were produced by double layer plaque assay using Shieh medium. Plates were incubated 1-2 days at RT, after which 5 ml of medium was applied on the confluent plate and incubated > 5 hours (8°C, shaking). Lysate was collected and filtered (0.2 µm) and stored at 8°C. Larger scale propagation of FLiP and FLiP-2 was done in liquid cultures in Erlenmeyer flasks as previously described for FLiP using lake-water based medium (J-Shieh) on top of a layer of solidified J-Shieh [2% agar (w/v)] (Mäkelä et al. 2024).

Drop titration on double layer agar assay was applied to determine lysate titers. Dilution series of phage lysates in 2 µl or 10 µl drops were pipetted on the plates. After incubating (2 days, RT) the PFU ml^-1^ was determined.

### Transmission electron microscopy

For transmission electron microscopy MaF61 filtrated lysate was applied on Formvar coated and glow-discharged copper grids for 60 seconds. Filtered (0.2 µm) 2 % phosphotungstic acid (PTA) was added on top of blot-dried samples for 10 seconds. After blotting and air-drying grids were imaged with JEOL JEM-1400HC TEM with 80 kV and RADIUS 2.1 (Build 20150, Emsis GmbH) software.

### Phage DNA extractions

Phage genomic DNA was extracted from 2 ml of filtered phage lysate using a zinc chloride precipitation method modified from Santos (M. A. Santos 1991). Incubation with DNase I and RNase A (final concentrations 0.001 mg ml^-1^ and 0.01 mg ml^-1^ respectively, 30 min, 37 °C) was followed by addition of zinc chloride (final concentration 40 mM) and incubation (5 min, 37 °C). Precipitate was centrifuged (1 min, 10 000 rpm) and the pellet was resuspended into 1 ml of TES buffer (0.1 M Tris-HCl pH8,0; 0.1 M EDTA; 0.3 % SDS) and incubated (15 min., 60 °C), followed by another incubation with proteinase K (final concentration 0.4 mg ml^-1^, 100 min., 37 °C). DNA was purified by continuing either with (1) phenol-chloroform method or (2) GeneJET kit:

(1) In phenol-chloroform method, 120 µl NaAc (3 M, pH 5.2) was added and incubated on ice (15 min). After centrifugation (1 min, 12 000 rpm), 1 volume of Phenol/CH_3_Cl/IAA (25:24:1) was added to the supernatant. The mixture was centrifuged (5 min, 8000 rpm) and the top phase containing DNA precipitated with 1 volume of ice-cold isopropanol and centrifuged (15 min, 12000 rpm), washed with ice cold ethanol (70 %) and centrifuged (10 min, 12000 rpm). Dried pellet was resuspended to 20 µl of Elution buffer from DNeasy Blood & Tissue kit (Qiagen).
(2) After proteinase treatment DNA was bound into purification columns of GeneJET Genomic DNA Purification Kit (Fermentas) using a mix of guanidine hydrochloride (final concentration 2 M) and EtOH (final concentration 25 %). Wash and elution steps were done according to manufacturer’s instructions. Elution volume of 100 µl was used. Purity and concentrations of extracted DNA were measured by Nanodrop and Qubit (High Sensitivity dsDNA Assay Kit).

### Genomic sequencing and sequence analysis

The genome of FLiP-2 was amplified by PCR (supporting information) to produce dsDNA. Genomes of FLIP-2 and MaF61 were commercially sequenced using Illumina NovaSeq and paired-end 150 bp sequencing (Novogene).

Illumina short reads were quality trimmed using BBduk, and 1 % of the trimmed reads were de novo assembled using Velvet (Zerbino and Birney 2008) with K-mer length 73 by using plug-in modules in Geneious Prime (Biomatters Ltd) to produce a draft genome. FLiP-2 genome was finalized by mapping all the short reads on the draft genome with Bowtie2 built in breseq (Deatherage and Barrick 2014) and manually correcting the genome sequence accordingly. The genomic differences between FLiP-2 and FLiP were determined by whole genome alignment and for each gene by pairwise alignments of homologous nucleotide and amino acid sequences using MUSCLE (Edgar 2004). A BlastP (Altschul et al. 1990) search was performed using FLiP MCP sequence against non-redundant protein sequence database during spring 2024. Surrounding sequences from the observed hits were inspected for other similarities with FLiP (including replication initiation protein, lytic enzyme and gene synteny). Genomic comparison of FLiP, FLiP-2, and related FLiP-like viral metagenomic assembly (vMAG) and putative prophage sequences was done with clinker (Gilchrist and Chooi 2021) which is based on all-against-all amino acid sequence similarity.

The assembled MaF61 genome contig (167 814 bp long with ∼8000 coverage) was run in July 2022 in NCBI BLASTn and together with the short reads in PhageTerm (Garneau et al. 2017) on the Galaxy@Pasteur server to determine the genome ends and packaging system. The contig was also checked with CRISPRCasFinder (Couvin et al. 2018) to recognize possible CRISPR arrays and associated Cas genes and tRNA finder to identify tRNAs. Glimmer (Delcher et al. 1999) and GeneMark (Besemer and Borodovsky 2005) softwares were used to predict open reading frames (ORFs). HMMER suite (Finn et al. 2011), InterPro (Hunter et al. 2009), BLASTp (Altschul et al. 1990) and PSI-BLAST (Bhagwat and Aravind 2007) were used to analyze the amino acid sequences translated from ORFs.

### Comparison of plaque morphologies of FLiP, FLiP-2 and MaF61

Double layer agar assay was used to detect possible differences in the plaque morphologies of FLiP, FLiP-2 and MaF61 at room temperature. Previously found shared hosts of FLiP and MaF61, *Flavobacterium* sp. B114, B167 and B330 (Mäkelä et al. 2024), were used on Shieh (Song et al. 1988) agar plates. After two days of incubation, plates were imaged with ChemiDoc MP Imaging system (Bio-Rad) using application for Ethidium Bromide Gel (602/50, UV Trans) and Auto Optimal exposure.

### Host ranges and efficiencies of plating of FLiP, FLiP-2 and MaF61

In determination of host ranges and efficiencies of plating (eop), Shieh medium at RT was used as the baseline condition for comparisons. *Flavobacterium* sp. strains B28, B80, B114, B167, B169 (Laanto et al. 2011), B205 (unpublished) and B330 (Laanto et al. 2017) were used in the first experiment (Experiment A) to determine the host range and eop of FLiP, FLiP-2 and MaF61 at three temperatures (RT, 18°C and 8°C) in Shieh. Plaques were counted after two or more days when no further bacterial growth or changes in phage infection were observed.

In experiment B, three other media were used in addition to Shieh: High-tryptone cytophaga (HTC; (Pate and De Jong 1990)), LB (Lennox) (NaCl concentration 5 g/L) and TYES (Holt et al. 1989). *Flavobacterium* sp. strains, which FLiP could infect in Experiment A, were used: B80, B114, B167 and B330. Plates were incubated at RT.

In the third experiment (Experiment C) eop of FLiP, FLiP-2 and MaF61 in LB (Lennox) (5 g/L NaCl) and in LB (Luria) (0,5 g/L NaCl) were compared to Shieh at RT using *Flavobacterium* sp. strains B114, B167 and B330 as hosts.

### Phage decay at room temperature

FLiP, FLiP-2 and MaF61 decay in Shieh and in filtered lake water was followed at RT. All samples were prepared in triplicates: 25 ml of Shieh medium or autoclaved lake water from Jyväsjärvi with 5 x 10^6^ PFU ml^-1^ phage was added into 50 ml tube and gently vortexed (1-2 seconds). Tubes were standing still between samplings and lightly vortexed (1-2 seconds) before samples were taken. The first sampling was done 1 hour after starting the experiment. Next samplings were done once a week during the first 5 months and every other week during the following ∼4 months. PFU ml^-1^ was determined on B330 as described above.

### Phage growth in low-oxygen environment

Importance of oxygen for host bacterial growth and FLiP infection was tested by incubating plates at RT both under normal and lowered concentration of oxygen. The low-oxygen environment was created by using Millipore Microbiology Anaerocult P reagent with Millipore Microbiology Anaerotest. However, the incubation cannot be considered as strictly anaerobic: It took ∼5-6 hours to reach anaerobic condition according to indicators. Either plain Shieh-soft (0,7 % agar) or supplementation with 5 mM or 10 mM NaNO_3_ as an oxidizing agent was used. Double layer plaque assay was applied in three ways (Figure 1) using host bacteria B114, B167 and B330 at RT: (A) 100 µl of phage dilution was mixed with 100 µl of overnight bacterial culture and 3 ml of Shieh soft. (B) Drops (10 µl) of FLiP, FLiP-2 and MaF61 lysate dilutions were plated on each host. Plates (A and B) were incubated in anaerobic bags (5 days). (C) First only the plated bacteria were subjected to low-oxygen condition for 5 days after which plates were retrieved from anaerobic bags and dilutions of phage lysates were immediately pipetted on them. In all cases, after low-oxygen treatment, plates were incubated in the presence of normal concentration of oxygen at RT for 2 days after which bacterium was well grown, and no more phage plaques appeared. Control plates were plated at the same time and incubated at RT in normal oxygen condition for 2 days.

**Figure 1.**
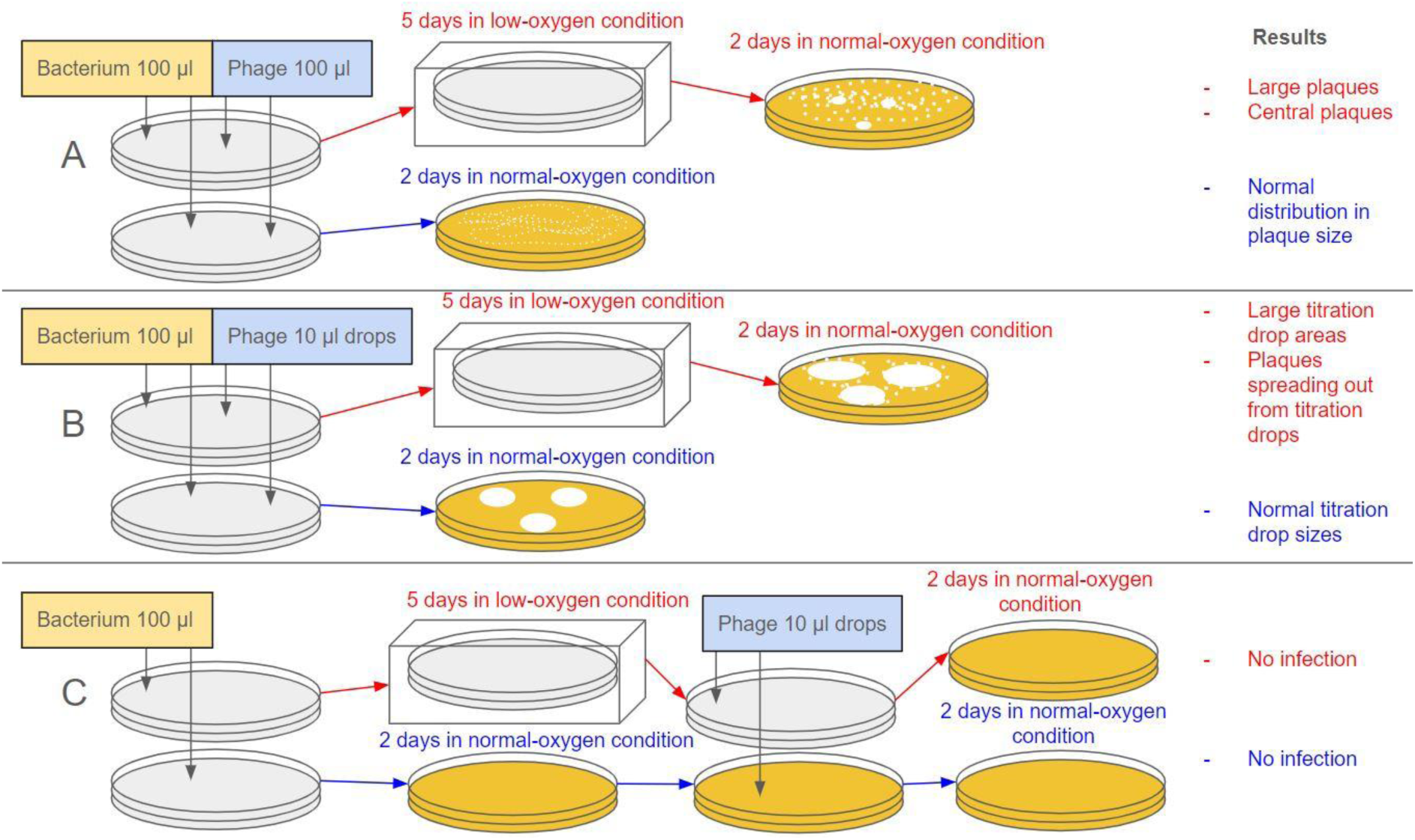
Titration plates incubated in low-oxygen or in normal-oxygen conditions. Plate color represents the growth of bacterium: White plate = No bacterial growth. Yellow plate = Full grown bacterial lawn. Text in red font describes methods and results of low-oxygen incubation and text in blue font describes those for normal-oxygen incubation.

### FLiP propagation, plaque morphology and host bacterium cell morphology in the presence of antibiotics

Different types of antibiotics with different modes of action, were used (ampicillin, kanamycin, tetracycline and tobramycin) to see the effect of each antibiotic on the growth of *Flavobacterium sp.* B114, B167 and B330. Antibiotics were added in Shieh medium to concentrations of 0, 25, 50 and 100 µg ml^-1^. Overnight grown cultures of bacteria (RT, 120 rpm) diluted to optical density (OD) A595 ∼0,3 were added (20 µl) to Shieh with and without antibiotics (180 µl), into a 96-well plate in four replicates. OD of the samples was measured at 595 nm using Tecan Spark microplate reader in the beginning of the experiment and after incubating 3, 4, 5, 24 and 49 hours at RT.

For a deeper analysis of the effect of ampicillin on the phage-host interactions, further experiments with strain B114 were conducted. The effect of ampicillin on phage titers (FLiP and FLiP-2) and on bacterial growth in concentrations of 0, 50, 100 and 500 µgml^-1^ with and without the presence of phages were studied in 96-well plates. Initial phage titer 1,0 x 10^5^ PFU ml^-1^ and bacterial cell density approximately 5,0 x 10^7^ CFU ml^-1^ were used. 178 µl of antibiotic dilution, 20 µl of bacterial culture and 2 µl of phage or Shieh (as control) were added in the wells in three replicates. OD A595 was measured using Multiskan FC Microplate Photometer (Thermo Fisher) and SkanIt Software for Microplate Readers (Thermo scientific) in the beginning of the experiment and once every 24 hours for three days. The effect of phage and antibiotic on bacterial growth was analysed with two-way ANOVA with multiple comparisons (LSD) using GrapPad Prism 10. In the end of the experiment, the phage titers were determined and microscopy imaging of cell morphologies with and without the presence of ampicillin and FLiP was performed. Bacterial cells were stained with Safranin after 24 hours of incubation for microscopy. Leica DM500 microscope with Leica ICC50 W camera were used for imaging cell morphologies using eyepiece field number 20 mm with 10x (FOV 2000 µm) and 40x (FOV 500 µm) objectives. Scale bars were added to images using ImageJ version 1.54i.

## Results

### FLiP-2 is closely related to FLiP

A new *Finnlakevirus* named FLiP-2 infecting *Flavobacterium* sp. B330 was isolated from Lake Jyväsjärvi in August 2021, almost exactly 11 years after FLiP. Sequencing of the FLiP-2 genome revealed 9176 nt long circular genome (accession number PV425005) with 34.5 % GC and 96,7 % nucleotide identity to FLiP. Consequently, according to ICTV standards, FLiP-2 is considered a strain of FLiP (Adriaenssens and Brister 2017).

The whole genome alignment of FLiP and FLiP-2 revealed 305 genetic differences, most of which were distributed among all the ORFs (Table 1). 26 of the nucleotide differences were located on non-coding regions. The most divergent genes were the non-structural genes at the beginning of the genome (Table 1; ORF 2 and ORF 3). Four of the 16 predicted genes were identical in amino acid sequence level: ORF 5, gp9, gp11 and gp12. Function of ORF 5 is unknown but gp9, gp11 and gp12 code for structural proteins, gp12 being the FLiP penton (Kjezar et al., 2022) (Figure 2; Table 1). Most of the genes (8) shared a 98-99 % amino acid level similarity; among them major capsid protein (MCP) (gp8), lytic transglycosylase (gp14) and replication initiation protein (ORF 15).

**Table 1.** Differences between Finnlakevirus strains FLiP and FLiP-2 genes and amino acid sequences.

| FLiP ORF / gp | Genome coordinates (start-stop) | Putative function | Length (bp) | Length (aa) | Differences (nt) | Differences (aa) | Identity (nt) | Similarity (aa) |
| --- | --- | --- | --- | --- | --- | --- | --- | --- |
| 1 | 155-445 |  | 291 | 97 | 3 | 1 | 99.0% | 99.0% |
| 2 | 540-782 |  | 243 | 81 | 16 | 6 | 93.4% | 92.6% |
| 3 | 836-1138 |  | 303 | 101 | 18 | 9 | 94.1% | 91.1% |
| 4 | 1135*-1557 | repressor | 423 | 141 | 25 | 7 | 94.0% | 95.7% |
| 5 | 1799-2008 |  | 210 | 70 | 6 | 0 | 97.1% | 100.0% |
| 6 | 2011-2229 |  | 219 | 73 | 5 | 2 | 97.7% | 97.3% |
| 7 | 2703-2957 | S, putative lytic enzyme, putative endolysin | 255 | 85 | 5 | 2 | 98.0% | 97.6% |
| 8 | 3036-3971 | S, MCP | 936 | 312 | 34 | 5 | 96.4% | 98.4% |
| 9 | 3992-4363 | S | 372 | 124 | 14 | 0 | 96.2% | 100.0% |
| 10 | 4385-4861 |  | 477 | 159 | 8 | 1 | 98.3% | 99.4% |
| 11 | 4845-5321 | S | 477 | 159 | 15 | 0 | 96.9% | 100.0% |
| 12 | 5324-5770 | S, penton | 447 | 149 | 14 | 0 | 96.9% | 100.0% |
| 13 | 5770-6780 | S, putative spike | 1011 | 337 | 27 | 4 | 97.3% | 98.8% |
| 14 | 6780-7445 | S, lytic transglycosylase | 666 | 222 | 28 | 2 | 95.8% | 99.1% |
| 15 | 7510-8853 | replication initiation protein | 1344 | 448 | 53 | 3 | 96.1% | 99.3% |
| 16 | 8865-9110 |  | 246 | 82 | 8 | 1 | 96.7% | 98.8% |

*S=structural protein; \* start of FLiP ORF 4 annotation updated (1138 -> 1135) in order to correspond FLiP-2 annotations.*

**Figure 2.**
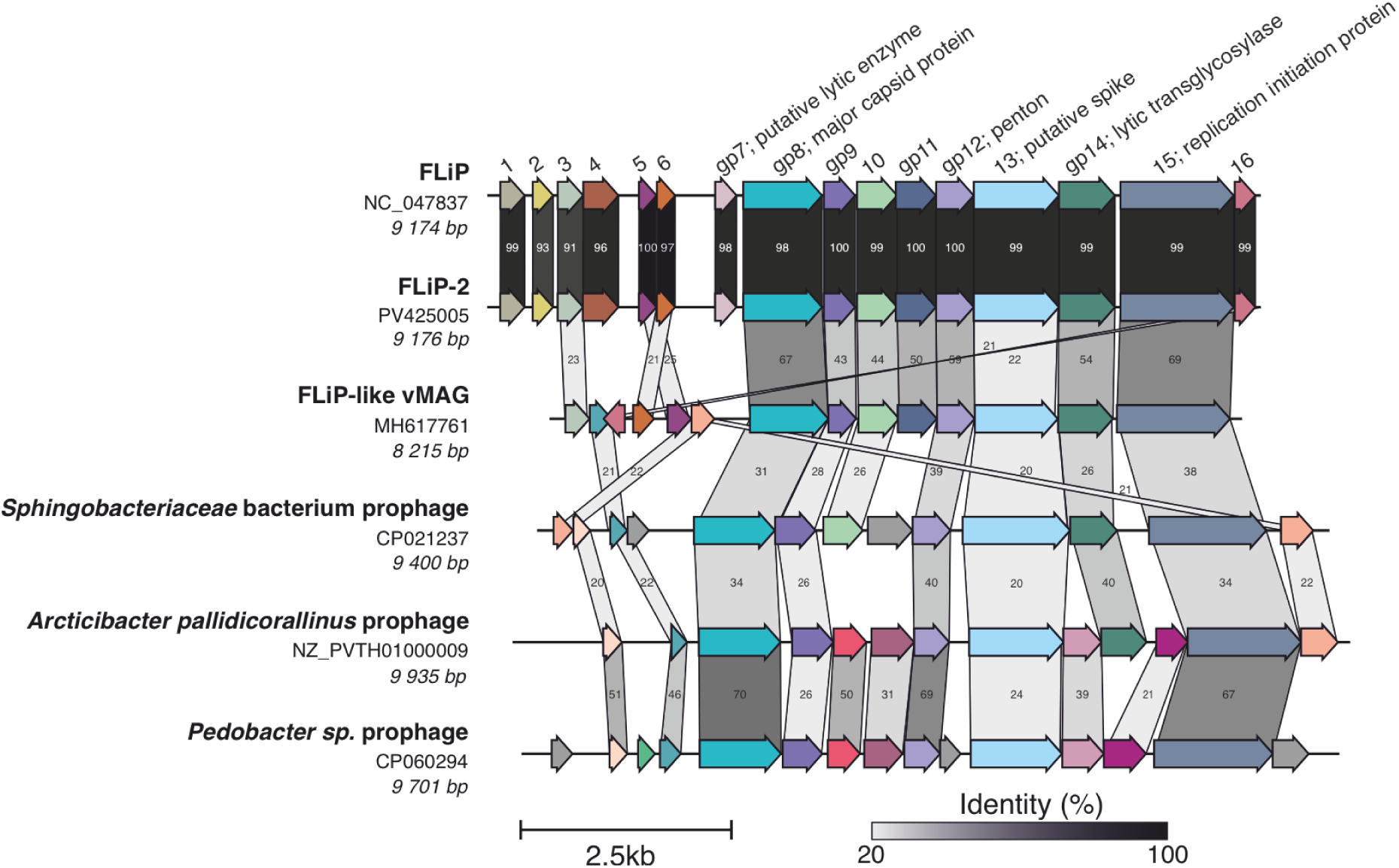
Genome comparison of FLiP, FLiP-2 and related FLiP-like viral metagenome-assembled genome (vMAG) and putative prophages found from bacterial genomes. ORF numbers refer to original FLiP annotations. Amino acid sequence similarity is presented as percentage between related ORFs/genes (threshold 20%).

A BLASTp search using FLiP MCP sequence resulted into hits to a metagenome assembled Bacteriophage sp. (65,92% identity, 100% query coverage) and three bacterial genomes belonging to the class *Sphingobacteriia* (*Pedobacter* 29,91% identity, 98% query coverage; *Sphingobacteriacae* 34,34 % identity, 99% query coverage; and *Arcticibacter*, 32.86 % identity, 83% query coverage), that are members of the same phylum as *Flavobacterium* species, *Bacteroidota*. Genes surrounding the hit were investigated revealing the overall genome synteny being similar to FLiP isolates (Figure 2). Indeed, genome alignment showed high levels of similarity especially among the structural genes and the replication initiation gene (Figure 2). While the function of the putatively non-structural genes located in the beginning of FLiP genome are left without function, an HHPred (Söding et al. 2005) search of ORF 4 in FLiP suggests homology with transcription regulators, especially to the DNA-binding domain of repressor protein in phage P22 (aa 80-136 in ORF 4; probability 95,24%; E-value 0.066). Overall, the gene synteny among the structural and replication genes seems to be conserved among FLiP isolates, viral MAG (vMAG) assembly and putative prophages, and the left side of the genome being more variable, containing putatively genes associated e.g. in gene regulation.

Also, another phage, named MaF61, that infects *Flavobacterium* sp. strains B114, B167 and B330 was isolated in November of 2021 from the same location as FLiP-2. Under TEM, MaF61 was seen to have a typical myovirus-like morphology with a capsid (∼110 nm facet to facet) and ∼170 nm long contractile tail (Supporting information Figure S1). Sequencing of MaF61 genome resulted in 167 814 bp long genome (accession number PQ858689) with a headful packaging mechanism and 195 ORFs were predicted. A putative function was identified only for 45 (23%) of the MaF61 ORFs (Supplementary file 2). MaF61 is so far the only known tailed dsDNA phage, which can infect three of the *Flavobacterium* hosts of FLiP (Mäkelä et al. 2024) making it a good control phage to study whether certain traits of FLiP and FLiP-2 are purely phage-dependent or if same phenomena can be seen with a dsDNA phage.

PCR results for the FLiP MCP gene were positive in the majority (23 out of 32) of water samples collected from sites other than where FLiP-2 and MaF61 were isolated (Supporting Information Table 1). In contrast, positive results for the FLiP Rep gene were less common, with only 6 out of 32 samples testing positive.

### FLiP and FLiP-2 plaque morphologies and sizes depend on the host

Plaque sizes and morphologies of FLiP, FLiP-2 and MaF61 were compared at RT on common hosts B114, B167 and B330. FLiP and FLiP-2 showed a distribution of both small and larger plaques on all hosts (Figure 3A) as observed for FLiP previously (Mäkelä et al. 2024), while MaF61 plaques were relatively uniform in size and mainly smaller than *Finnlakevirus FLiP* plaques (Figure 3B). The largest FLiP plaques were observed on B114, while FLiP-2 plaques were remarkably smaller on this host. On B167, FLiP-2 plaques were slightly larger than FLiP plaques. The opposite was observed on B330. Both clear and turbid FLiP and FLiP-2 plaques were observed, depending on the phage-host combination: Distribution of clear and turbid plaques produced by both phages were seen on B167 (Figure 3A). While FLiP plaques on B114 were clear, FLiP-2 plaques were turbid. FLiP plaques on B330 were either clear or turbid, but only turbid FLiP-2 plaques were observed on B330. Most *Finnlakeviridae* plaques had at least to some extent fuzzy edges whereas MaF61 plaques were clear and had sharp edges.

**Figure 3.**
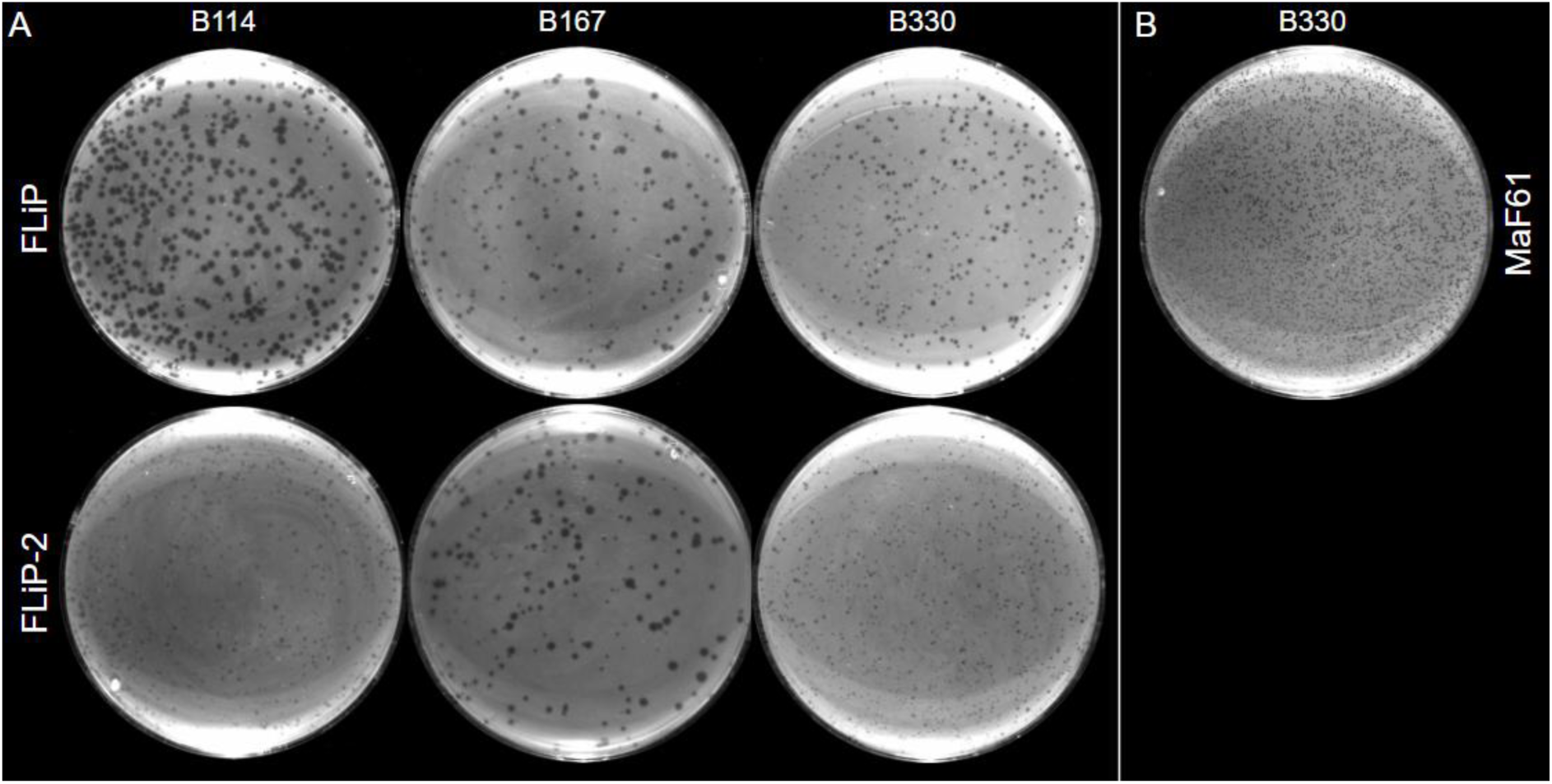
FLiP, FLiP-2 and MaF61 plaque sizes and morphologies in double layer agar assay. A) Plaque size and morphology of FLiP and FLiP-2 at room temperature on Flavobacterium sp. B114, B167 and B330 on Shieh agar. B) Plaque size and morphology of MaF61 at room temperature on Flavobacterium sp. B330 on Shieh agar. Plates were imaged with ChemiDoc MP Imaging system (Bio-Rad).

### Host ranges and efficiencies of plating of FLiP and FLiP-2 vary differently depending on nutrient availability and temperature

The host range and efficiency of plating (eop) of FLiP, FLiP-2 and MaF61 on *Flavobacterium* sp. strains at temperatures 8℃, 18℃ and RT (∼21℃) were tested. FLiP was able to infect 4 *Flavobacterium* strains: B330, B167, B114 and B80, consistent with what has been observed in previous host range experiments (Mäkelä et al. 2024). FLiP-2 and MaF61 could infect the same strains except for B80, in which they caused growth inhibition but not plaques. Eop of FLiP was lower with B80 compared to other hosts (Figure 4A-B), and FLiP lost its ability to infect B80 in further experiments. FLiP and MaF61 also occasionally lost their ability to infect B167 or B114 even without changes in culture conditions. Inconsistent infection patterns have not been observed with FLiP-2 thus far. MaF61 could infect also additional bacterial strains, B28, B169 and B205. Temperature did not affect phage infection with strains B167 and B330 (Figure 4A). While infection of all phages on B114 was efficient both at RT and at 18℃, none of the phages could form plaques on B114 lawn at 8℃: Only FLiP could lyse B114 at cold temperature observed as clear titration drop area on the bacterial lawn, while FLiP-2 and MaF61 could only inhibit bacterial growth.

**Figure 4.**
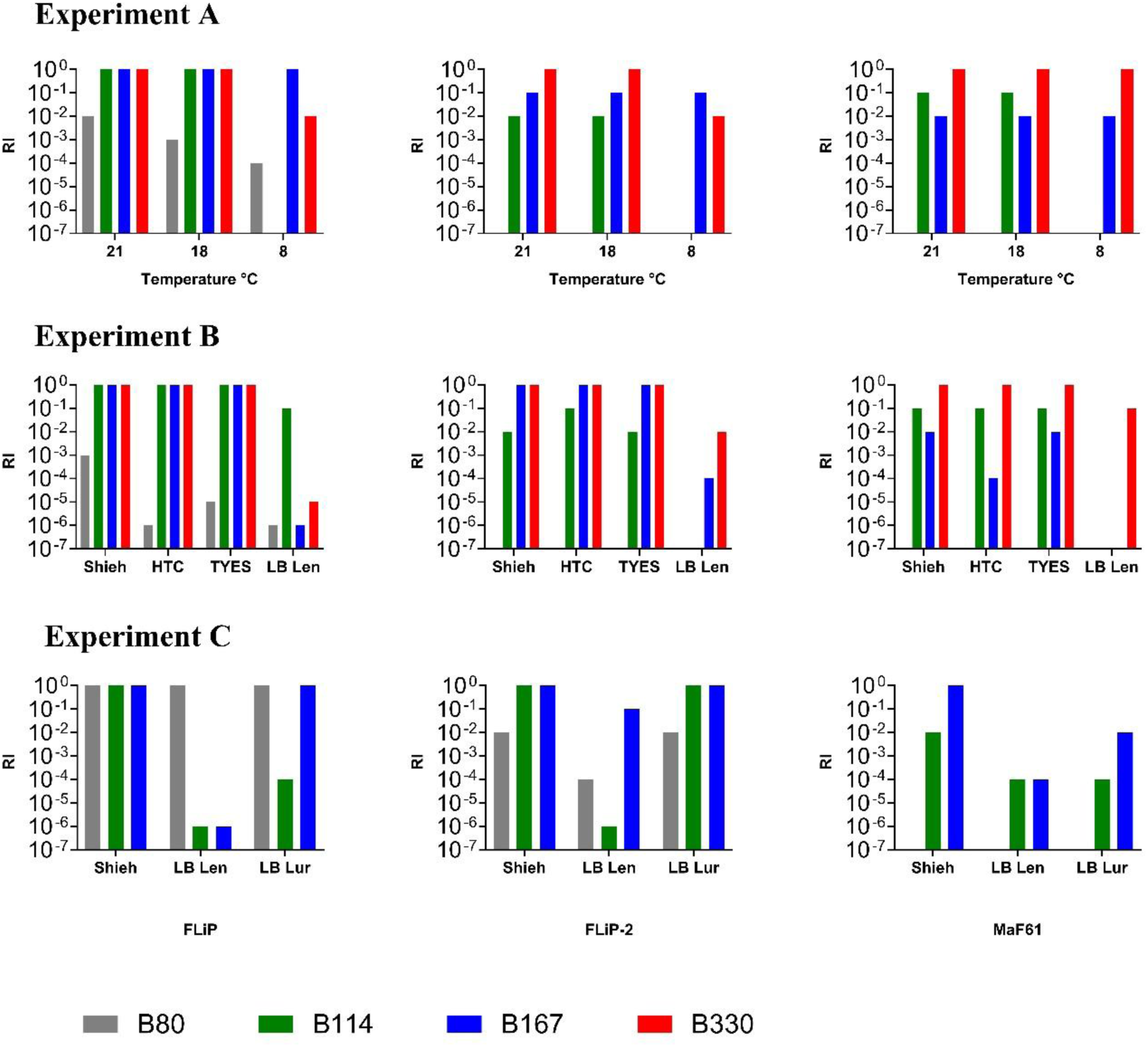
FLiP, FLiP-2 and MaF61 relative efficiencies of infection (RI) compared to baseline conditions (Shieh, Flavobacterium sp. B330, room temperature) A) at room temperature (21℃), 18℃ and 8℃ in Shieh, B) at room temperature in Shieh, HTC, TYES and LB growth media and C) at room temperature in LB with NaCl concentrations of either 5 g (LB Lennox = LB Len) or 0,5 g/L (LB Luria = LB Lur) compared to Shieh.

Also, the effect of different growth media on eop was tested (Experiment B) using four different media (Shieh, HTC, TYES and LB) at RT. Only small differences between titers were observed when either Shieh, HTC or TYES were used (Figure 4B). Eop of all three phages on B167 was lower in LB compared to other media (Figure 4B). On B114, the difference was drastic between the phages: FLiP-2 and MaF61 only inhibited the growth, while the eop of FLiP was almost the same as with other media. The exact opposite was observed with B330; FLiP titer was very low, while FLiP-2 titer was lowered by only two orders of magnitude compared to other media (Figure 4B).

To further explore whether the 5g/L NaCl concentration in LB was the reason for the observed differences in phage infectivity, the concentration was lowered to 0,5 g/L. This increased titer of all phages, but the host-specific variation in this increase was substantial. (Figure 4C). MaF61 did not infect B114 in this experiment.

### FLiP and FLiP-2 remained infective for 50-70 days at room temperature

The decay of lipid-containing ssDNA phages FLiP and FLiP-2 at RT was fast compared to dsDNA phage MaF61, both in Shieh medium and in lake water. Infective FLiP particles were found for 50 days in lake water and for 70 days in Shieh (Figure 5), whereas infective FLiP-2 particles were found for 70 and 63 days, respectively. MaF61 decay was much slower. Differences between Shieh and lake water could be seen after ∼70 days: While ∼104 PFU ml^-1^ of MaF61 particles were found until the end of the experiment (266 days) in lake water, only ∼102 PFUml^-1^ of MaF61 were left in one of the three replicates in Shieh.

**Figure 5.**
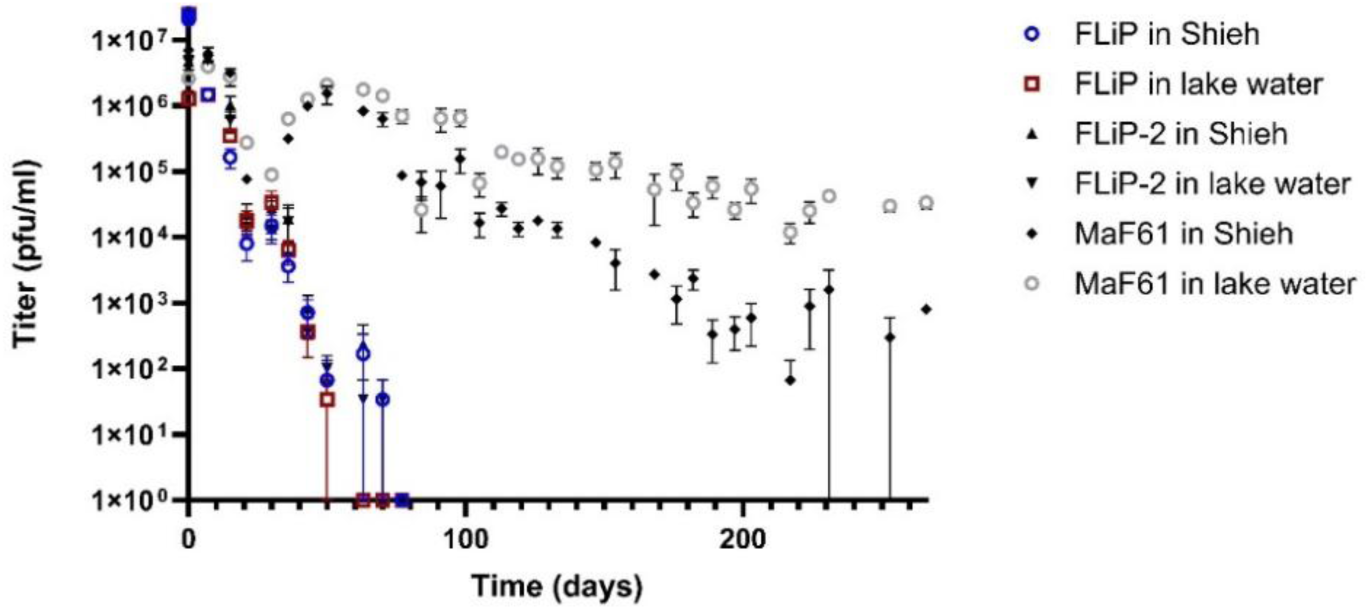
Phage decay at room temperature (mean +/- SEM of 3 replicates). Titer of all phages was 2,50 x 10^7^ PFU ml^-1^ in the beginning of the experiment.

### Low-oxygen incubation leads to enlarged lysis area and spreading of plaques from the original infection site

In a preliminary experiment (data not shown), presence of NaNO_3_ did not affect bacterial growth or phage infection in any way in low-oxygen or normal-oxygen conditions and was not used in further experiments. Furthermore, preliminary experiments showed that FLiP and FLiP-2 interactions with their hosts changed similarly under low-oxygen conditions: The results were comparable in terms of phage titers, and both strains exhibited enlarged lysis areas as a result of anaerobic incubation. Therefore, only the results for FLiP and MaF61 are presented below.

Bacteria did not grow in low-oxygen conditions during 5-day incubation but formed a lawn in two days after retrieving to normal-oxygen environment (Figure 1A-B). Two days were also needed for full grown bacterial lawn in normal-oxygen condition without low-oxygen phase (Figure 1A-B). Although the bacteria did not grow during the low-oxygen incubation, they lost their susceptibility to FLiP (Figure 1C).

Plaque sizes and clear areas caused by phage lysis in drop titration (lysis area) were first surveyed for FLiP in normal and in low-oxygen conditions. Lysis areas caused by FLiP were clearly larger when incubation included a low-oxygen phase (Figure 6). In addition, spreading of plaques from the lysis area edges to the surroundings could be seen when incubation contained a low-oxygen phase. This phenomenon was most clearly seen after low-oxygen incubation of B114 which caused some of the FLiP plaques to form “central plaques” from which more plaques were spread to the surrounding area (Figures 1A and 6B). To compare the effects of lowered oxygen concentration between FLiP and MaF61, two replicate experiments were conducted. In each of the experiments both phages were added on the same plate to ensure equal oxygen concentration. FLiP lysis area size increased more in anaerobic incubation compared to the lysis area size of MaF61 (Figure 6C). The titer of MaF61 decreased significantly following low-oxygen incubation with both B167 and B114 hosts but remained almost unchanged with the B330 host. Titer of FLiP was not much affected by low-oxygen incubation (Figure 6C).

**Figure 6.**
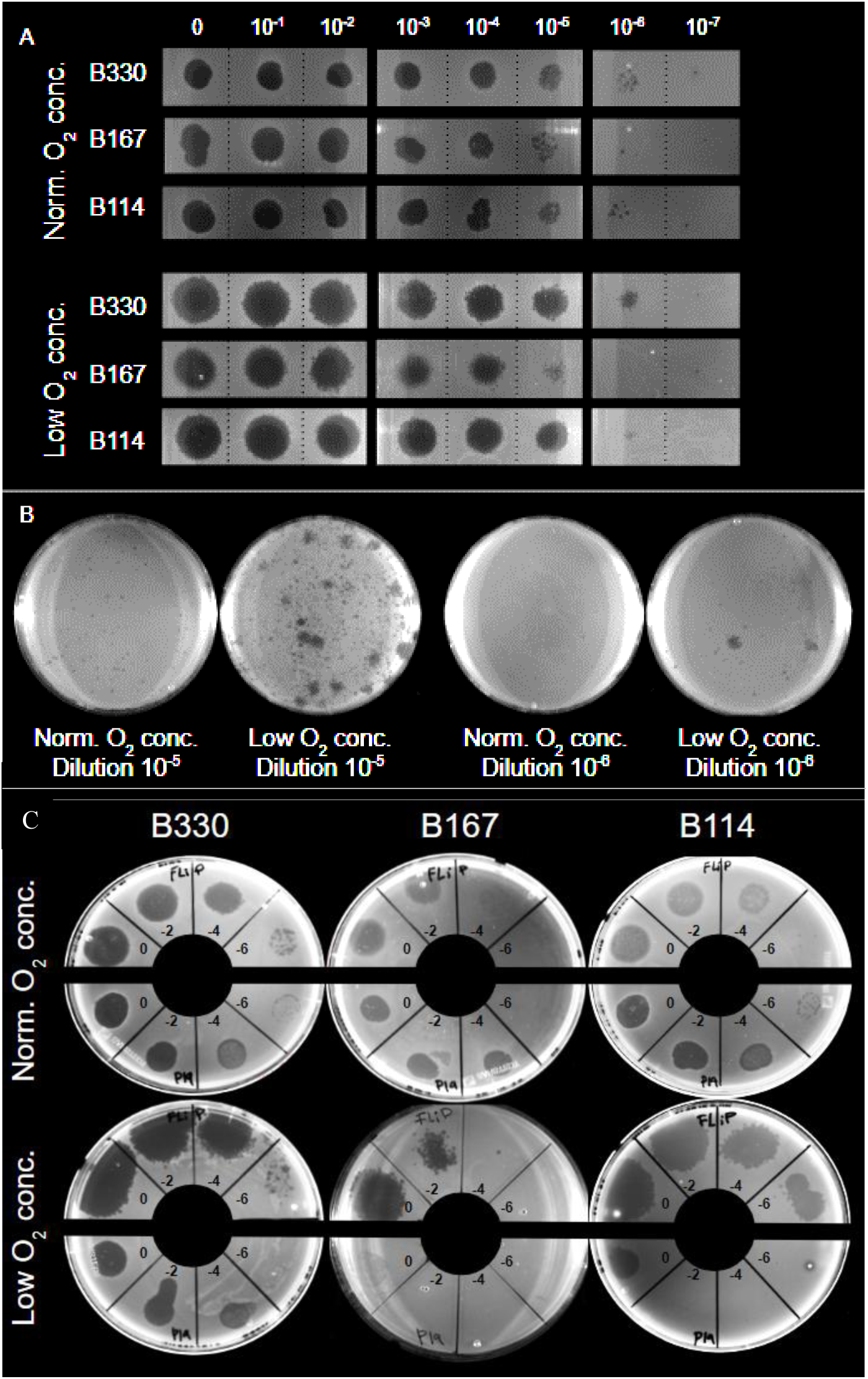
FLiP lysis area sizes and plaque morphologies in double layer agar assay. Plates were either incubated two days at room temperature in presence of normal concentration of oxygen (Norm. O_2_ conc.) or 5 days in low concentration of oxygen followed by an additional two days at room temperature in normal oxygen concentration (Low O_2_ conc). A) FLiP plaque and lysis area morphologies on Flavobacterium sp. B330, B167 and B114. Undiluted FLiP lysate and a dilution series (10 µl each) were pipetted on plates. Plates imaged with ChemiDoc MP Imaging system (Bio-Rad). Figure composed of separate parts of plate images: Solid lines indicate edges of one part and dotted lines confine areas of each titration drop. B) 100 µl of either 1 x 10-5 or 1 x 10-6 dilution of FLiP lysate plated with Flavobacterium sp. B114. C) FLiP (on upper half of the plate) and MaF61(on lower half of the plate; P19 is the isolate name of the phage) dilutions in 10 µl drops on Flavobacterium sp. B330, B167 and B114.

### FLiP replication is efficient in the presence of antibiotics

Antibiotics were used to see how bacterial growth and phage-host interactions are affected by bacteriostatic substances. Bacterial strains B114, B167 and B330 had different sensitivities to each antibiotic type as detected in differences in growth in the presence of either ampicillin, kanamycin, tetracycline or tobramycin compared to antibiotic-free controls (Supporting information Figure S2). Bacterial cell morphologies were also affected by antibiotic treatments. Cell filamentation could be seen in all bacterial strains after treatment with any of the antibiotics (except for B167 and B330 after ampicillin treatment, which killed all the cells), and the elongation effect was strongest in B114 cells. B114 cells with both normal (∼2-10 µm) and significantly elongated length (even ∼100-200 µm) were detected in ampicillin treatments (Figure 7A). Some of the filamented cells did not show signs of division, but some had undergone division, at least partially, resulting in shorter cells, which still formed a cohesive cell chain (Figure 7A). Percentages of elongated versus normal-sized cells were similar in treatments with only antibiotic and in treatments containing both antibiotic and FLiP, but cell length increased even more in the presence of phage leading to cells even ∼300 µm long (Figure 7A).

**Figure 7.**
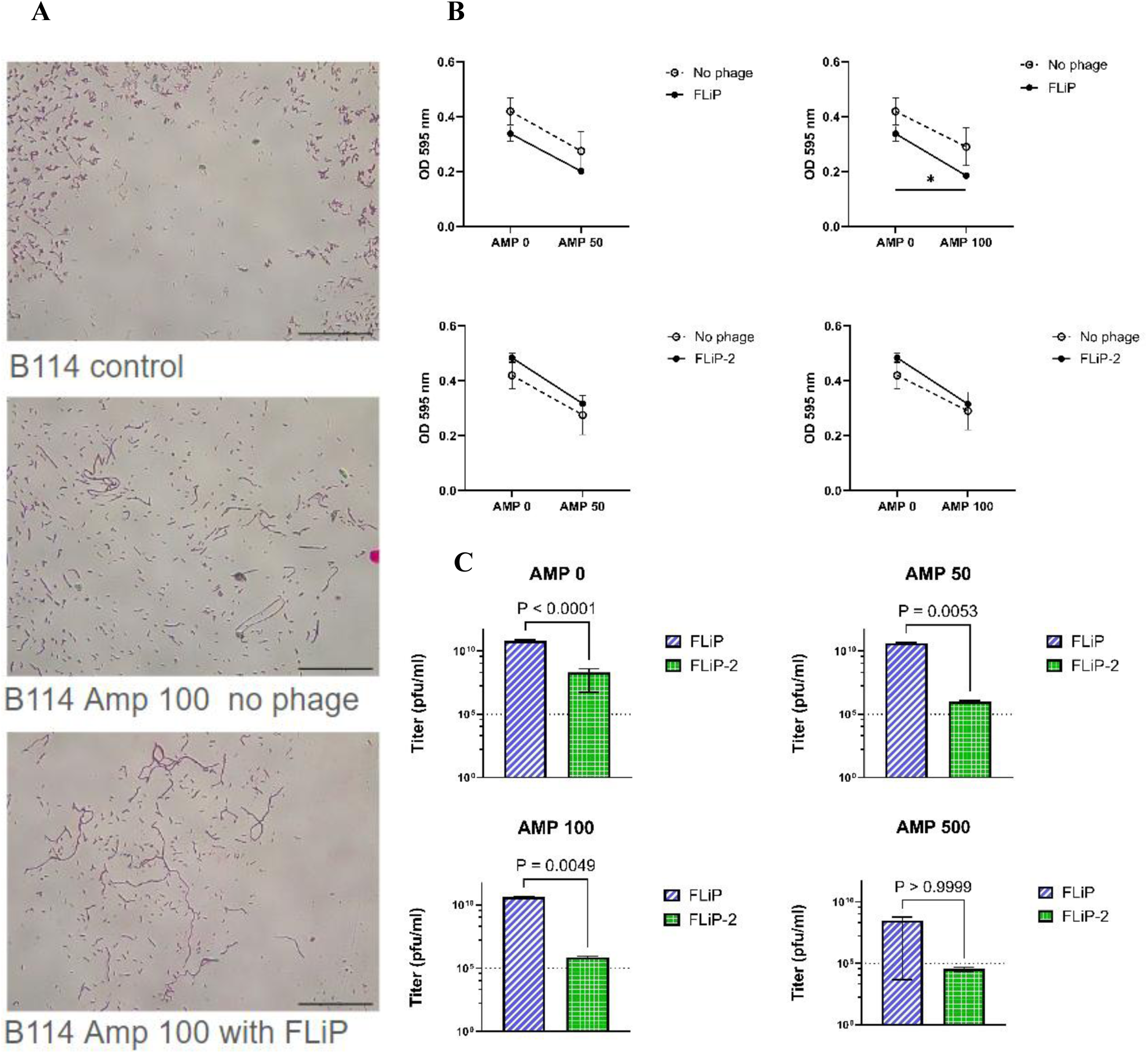
Growth of Flavobacterium sp. B114 and phages FLiP and FLiP-2 in the presence of ampicillin. A) Cell morphologies after 24 hours of growth in Shieh medium at room temperature. Scale bar in all images is 100 µm. B) Effect of ampicillin and phage on growth of B114 measured as optical density at 595 nm (mean +/- SEM of 3 replicates). C) Titers of phages after 3 days (mean +/- SEM of 3 replicates). Amp 0: Control without ampicillin. Amp 50: ampicillin concentration of 50 µg ml^-1^. Amp 100: ampicillin concentration of 100 µg ml^-1^. Amp 500: ampicillin concentration of 500 µg ml^-1^.

To analyse the possible synergistic effects of FLiP and ampicillin, we analysed their effects on bacterial growth using two-way ANOVA with multiple comparisons (LSD). The presence of ampicillin (50 µg ml^-1^) significantly reduced the bacterial growth (F1,8=9.39, p=0.016) whereas phage did not (Figure 7B). In the experiment with higher antibiotic concentration (ampicillin 100 ug ml^-1^) a similar result was obtained for ampicillin (F1,8=9.39, p=0.013). However, addition of ampicillin caused a significant decrease in bacterial growth in the presence of FLiP (LSD, p=0.042), indicating a possibility of phage-antibiotic synergy.

Similarly, antibiotics had a significant impact on bacterial growth with experiments done with FLiP-2 (amp50: F_1,8_=12.3, p=0.008; amp100: F_1,8_=11.7, p=0.009). Here, however, the bacterial growth in presence of FLiP-2 was higher than in the control treatment (Figure 5B), but interestingly in both antibiotic concentrations multiple comparisons revealed that ampicillin caused a significant decrease in bacterial growth in the presence of FLiP-2 (LSD, p=0.029 for ampicillin 50 µg ml^-1^ and 0.026 for ampicillin 100 ug ml^-1^).

FLiP titer increased in all ampicillin treatments (0-100 µg ml^-1^) to the level of 10^10^ PFU ml^-1^. FLiP titer even increased in one of the replicates with 500 µg ml^-1^ concentration to the level of 10^8^ PFU ml^-1^. FLiP-2 replication was weaker than FLiP in controls without ampicillin reaching barely the level of 10^8^ PFU ml^-1^. Even the lowest concentration of ampicillin prevented FLiP-2 replication almost completely (Figure 7C).

## Discussion

ssDNA phages have been shown to be prevalent in aquatic microbial communities (Tucker et al. 2011; Székely and Breitbart 2016). Due to the relatively small number of ssDNA phage isolates studied so far, our current understanding on their life cycles likely represents only a glimpse of the overall picture. While more research is needed for definitive conclusions, the existing evidence suggests that ssDNA phages likely possess a diverse array of life cycle strategies and ecological impacts under different conditions (Nguyen et al. 2023; Székely and Breitbart 2016; Tucker et al. 2011). This hypothesis, though compelling, awaits further confirmation through more extensive sampling and analysis of ssDNA phages across various environments. Here, we present a new isolate of the ssDNA *Finnlakevirus FLiP*, named FLiP-2. This strain of *Finnlakevirus FLiP* was isolated from the same location (Lake Jyväsjärvi, Central Finland) 11 years after the original isolation of FLiP, confirming that *Finnlakeviridae* has established prevalence and is an active member of the microbial community in this lake. In this study, we also identified FLiP-like sequences from a metagenomic assembly and bacterial genomes. In addition, previous findings that homologs of MCP of both FLiP and structurally similar ssDNA phage phiCjT23 along with their replication (Rep) protein are found from aquatic metagenomic contigs and genomes among Bacteroidota hosts suggests that the prevalence of this type of phages is geographically broad (Kjezar et al., 2022; Yutin et al. 2018). Our PCR screening of freshwater samples in Finland revealed that sequences similar to FLiP MCP gene were commonly detected, while sequences similar to FLiP Rep gene were less abundant. These findings suggest the presence of phages with similar capsid structures but possibly divergent replication initiation mechanisms. This observation underscores the importance of isolating and characterizing a broader range of phages to elucidate the true diversity within the currently unexplored viral taxa.

Previous studies of phage FLiP have not defined its lytic nature (Mäkelä et al., 2024; Laanto et al., 2017). Both phages, FLiP and FLiP-2, formed clear plaques on all hosts showing lytic potential while the occasional inability to infect two of the three host strains could indicate alternative life cycle to lytic infection (Mäkelä et al., 2024). Whether the hit to the ORF04 suggests any function related to temperate life cycle remains to be elucidated. Moreover, previous and current findings of ssDNA phage FLiP, phiCjT23 and Cellulophaga phage phi48:2 -like sequences from genomes of bacteria in phylum Bacteroidota (Kjezar et al., 2022; Yutin et al., 2018; Holmfeldt et al., 2013) suggest these phages could exist as a prevalent prophage. Cellulophaga phage phi48:2 has also been shown to encode homolog of FLiP DJR MCP (Yutin et al., 2018) while the presence of a lipid membrane has not been identified, as has been for the other isolates.

In recent years, there has been a shift towards studying phage-bacteria interactions as ecological phenomena influenced by both abiotic and biotic environmental factors (Stone et al. 2019). These interactions are not necessarily governed by static patterns but rather demonstrate flexibility in response to changing conditions (Mäkelä et al. 2024; Stone et al. 2019; Gaborieau et al. 2023; Mäntynen et al. 2021). The current study was built on our previous exploration of FLiP life cycle characteristics, which revealed strong condition dependency of FLiP-host interactions (Mäkelä, Laanto, and Sundberg 2024). Here, we included both abiotic and biotic variations typical for boreal lakes to investigate the ecology of these interactions. Three phages, FLiP, FLiP-2 and MaF61, were used with three *Flavobacterium* hosts, representing the biotic diversity in boreal lake ecosystems. We examined how these phage-host interactions are influenced by abiotic conditions and stress factors typical of boreal lakes, such as temperature fluctuations, nutrient composition, oxygen availability, and the presence of antibiotics.

Experiments done in reduced oxygen levels enhanced our understanding of certain steps in the course of FLiP infection. First of all, low-oxygen incubation experiments gave further evidence that FLiP infection may depend on the initial attachment process to the host cell surface, consistent with our previous observations (Mäkelä et al. 2024). If only the host bacteria were exposed to low-oxygen conditions, they lost their susceptibility to FLiP infection. Although visible bacterial growth started only after recovery of normal oxygen levels, the bacteria were no longer susceptible to FLiP infection. In contrast, when both bacteria and phage were added immediately before low-oxygen incubation, bacterial growth also commenced only after the return to normal oxygen levels, but in this case, FLiP infection was successful. This suggests that the phage infection occurred either before or during the low-oxygen incubation period, which would mean that the window for successful FLiP infection may be closely tied to the initial attachment process of the host, potentially more likely than its growth phase. However, other factors could also influence this process. For instance, the cellular state during low-oxygen conditions, such as reduced metabolic activity or altered expression of surface receptors, might affect the susceptibility of host cells to FLiP infection. Further experiments are needed to elucidate these potential influences and to draw definitive conclusions about the mechanisms involved.

Another aspect of FLiP life cycle revealed during low-oxygen incubation experiments was the observation of enhanced lysis and migrating plaques, both phenomena also observed when incubated in other stressful conditions. FLiP plaques and lysis areas were very large and clear when the host had suffered from low-oxygen conditions, and there were plaques observed up to several millimeters around the original infection site, leaving an intact bacterial lawn in between. No plaques were seen on bacterial lawn when cells were not exposed to FLiP before low-oxygen incubation, ruling out the possibility of prophage induction caused solely by anoxic stress. A possible explanation for remote plaques could be a prolonged latency period or lysogeny in low-oxygen conditions, during which the infected bacteria migrate away from the original infection site, but the active infection with lysis starts only at normal oxygen level. Another possible cause could be bacterial cell filamentation, which in stressful conditions is known to lead to enlarged plaque size in several bacterial species such as *Escherichia coli, Salmonella enterica, Pseudomonas fluorescens and Staphylococcus lentus* (S. B. Santos et al. 2009; Kim et al. 2018; Comeau et al. 2007). Our current microscopy images revealed that the stress caused by combination of ampicillin and FLiP induced extreme cell elongation in *Flavobacterium* sp. B114 (Figure 6A). While these images suggest the formation of cells potentially hundreds of times longer than normal, more advanced microscopy techniques with higher magnification are necessary to precisely measure cell lengths and analyze morphologies in greater detail. This future analysis will provide a more accurate rendition of the observed filamentation effect, but effects of filamentation can already be discussed. In case only a single copy of the phage genome resides at the extreme end of a long filamented cell, a remote plaque could form after post-stress cell division. The latent presence of the phage in a cell that both filaments and moves vigorously could therefore explain the plaques far away from the original infection site.

One of the main aims of this study was to compare host interactions of FLiP and FLiP-2 using dsDNA phage MaF61 (myophage) as a control. Despite their high genome similarity, the plaque morphologies, life cycles and host ranges of FLiP and FLiP-2 were different, indicating that even small genetic differences may substantially affect phage characteristics. It is not known if FLiP is an ancestor of FLiP-2 or if both phages coexist in lake Jyväsjärvi. In the case of coexistence, the genetic differences between them could reflect different ecological niches or selective pressures. For example, one strain might have adapted to infect a different host or microenvironment in the same lake, leading to different selective pressures on certain genes. Notably, all known structural proteins had no or only a few amino acid changes, underscoring their essential roles and evolutionary constraint. However, the putative spike gene exhibited four amino acid substitutions. Even small changes in key residues of viral receptor binding proteins can lead to substantial functional consequences in the efficiency of host recognition and binding (Obadan et al. 2019; Peka and Balatsky 2023), which has been shown to alter phage-host interactions such as the rate of adsorption and the host ranges of phages (Jia et al. 2023). Thus, the observed amino acid differences in spike protein could impact host receptor binding and phage adsorption kinetics, potentially partly explaining the differences in infection patterns between FLiP and FLiP-2. Higher variation in sequence similarity was found in ORFs with unknown functions encoding non-structural proteins, suggesting these differences may play crucial roles in host interactions within the cell and in regulating infection processes, potentially further explaining the observed variations in plaque morphologies and efficiencies of infection.

Surprisingly, *Finnlakevirus FLiP* infections demonstrated at least similar flexibility under varying conditions compared to the tailed dsDNA phage MaF61, with 18 times larger genome size. In many cases, FLiP was able to infect its host bacteria in stressful conditions as efficiently as in more optimal conditions. The greatest difference in stress tolerance between different phage types was seen in anoxic conditions, as MaF61 infection efficiency markedly suffered with both B114 and B167 hosts, while FLiP reached normal titers and improved lysis on all hosts. Infection efficiency of FLiP-2 also suffered under some stressful conditions, such as in the presence of ampicillin, while FLiP infection was not much affected unless the ampicillin concentration was very high. Observed differences among phage species and strains suggest that bacterial stress responses alone cannot explain altered phage-host interactions under stress conditions. Instead, phage characteristics significantly influence the outcome.

*Finnlakeviridae* consistently produced larger plaques and lysis areas compared to the dsDNA phage MaF61. This may stem from differences in adsorption rates, lysis times, and virion size and morphology between MaF61 and FLiP (Gallet et al. 2011). Yet, the observed variation in plaque sizes among *Finnlakevirus FLiP* strains suggest that their characteristically large plaque size cannot be attributed solely to their small virion size and easily diffusing shape. Furthermore, FLiP plaque morphology and size changed when conditions changed. For instance, interactions in each phage-bacterium pair were differently affected by different growth media. These effects extended from plaque morphologies to efficiencies of infection. Whereas FLiP showed rather consistent infectivity with different hosts and media, FLiP-2 infection efficiency was 10 to 100 -fold weaker in certain growth media or host (especially B114). High concentration of NaCl was one key factor to inhibit infection of both *Finnlakeviridae* and MaF61, but the exact reason was not examined. Electrostatic interactions have been shown to affect phage abilities to bind to their receptors, and osmotic stress may, on the other hand, even inactivate phages (Moldovan et al. 2007; Puck et al. 1951; Ranveer et al. 2024). High salinity is atypical in freshwater environments and may induce bacterial stress responses at the molecular level, even if not apparent in growth patterns, which were not observed here in plate cultures.

Our research revealed distinct survival strategies among phages under stressful conditions. The tailed dsDNA *Flavobacterium phage MaF61* demonstrated remarkable particle stability, remaining infectious for several months at room temperature. This allows MaF61 to persist as an infective particle form, waiting for favorable conditions before initiating infection—a characteristic observed in some other dsDNA phages as well (Xu et al. 2023). In contrast, *Finnlakeviridae*, as lipid membrane-containing phage and thus more sensitive to external conditions, exhibit a different strategy. These phages may require more frequent host interactions in nature, particularly during warmer seasons, as their particles remain infective for only a few months. This aligns with previous findings that lipid-containing icosahedral dsDNA phages generally show lower particle stability compared to tailed phages (Ackermann, Tremblay, and Moineau, 2004). While the stability of FLiP outside the host is compromised, it has an excellent ability to preserve effective infection during or immediately after stress exposure. The different phage-host dynamics of a tailed dsDNA phage and a ssDNA phage with the same hosts highlight two divergent approaches for enduring unfavorable environments, and the contrast in survival tactics underscores the diverse adaptations phages have evolved to navigate the challenging ecological landscapes.

## Conclusions

Despite their relatively small genomes, phage life cycles demonstrate a degree of adaptability to environmental conditions. *Finnlakevirus FLiP* exhibits remarkable plasticity in its infection dynamics, with the timing and outcome of phage-host interactions being contingent upon a complex interplay of viral strain characteristics, host specificity, and environmental conditions. The observed disparities in host interactions between FLiP and FLiP-2 underscore the significance of even minor genetic variations within the *Finnlakevirus* genus. The capability of *Finnlakevirus FLiP* in stressful conditions to temporal flexibility in the lytic cycle suggests sophisticated mechanisms for navigating environmental stressors. Such adaptability may confer a significant ecological advantage, allowing Finnlakeviruses to maintain their presence in fluctuating aquatic environments while synchronizing their replication with favorable host and environmental conditions. Further research is crucial to fully elucidate the complex dynamics of *Finnlakevirus FLiP* infections. As phage responses to host status are diverse and the underlying mechanisms may be regulated by either the phage itself or the host, it is imperative to explore the genetic differences in the host bacteria as well as to determine the mechanisms of prolonged phage presence inside the hosts.

## Supporting information

Supplementary file 2 - ORFs of MaF61

## Acknowledgements

We want to thank Annukka Enervi, Svitlana Bublyk, Amanda Mäkinen and Noora Rantanen for assistance in the laboratory and Petri Papponen for technical assistance in the laboratory. We thank Jonna Kuha, Juha Ahonen and Ahti Karusalmi for collecting water samples from Lake Jyväsjärvi. The research was funded by research grants from Emil Aaltonen Foundation (#200260 L.-R.S.), Research Council of Finland (#346772, L.-R.S. and #354982, R.P.), Olvi Foundation (#201910409 K.M.) and The Finnish Concordia Fund (#20200077 K.M.). This project has received funding from the European Research Council (ERC) under the European Union’s Horizon Europe research and innovation programme (grant agreement No 101117204) (E.L.).

