## Supplementary file 2 - ORFs of MaF61 for "New Finnlakevirus isolate FLiP-2 provides insight into the ecology of ssDNA phages in Flavobacterium hosts"

| ORF name | Start nucleotide | End nucleotide | Nucleotide Length | Direction | Final annotation |
|---|---|---|---|---|---|
| ORF1 | 315 | 1751 | 1437 | reverse | |
| ORF2 | 2026 | 2301 | 276 | reverse | |
| ORF3 | 2294 | 2677 | 384 | reverse | |
| ORF4 | 2677 | 2889 | 213 | reverse | |
| ORF5 | 2897 | 3757 | 861 | reverse | |
| ORF6 | 3934 | 4176 | 243 | reverse | |
| ORF7 | 4193 | 4492 | 300 | reverse | |
| ORF8 | 4496 | 4789 | 294 | reverse | |
| ORF9 | 4797 | 5054 | 258 | reverse | |
| ORF10 | 5065 | 5343 | 279 | reverse | |
| ORF11 | 5345 | 5608 | 264 | reverse | |
| ORF12 | 5608 | 6369 | 762 | reverse | |
| ORF13 | 6436 | 6630 | 195 | reverse | |
| ORF14 | 6633 | 6935 | 303 | reverse | |
| ORF15 | 6925 | 7179 | 255 | reverse | |
| ORF16 | 7182 | 7718 | 537 | reverse | |
| ORF17 | 7708 | 9531 | 1824 | reverse | |
| ORF18 | 9873 | 10094 | 222 | reverse | |
| ORF19 | 10091 | 10420 | 330 | reverse | |
| ORF20 | 10469 | 10615 | 147 | reverse | |
| ORF21 | 10666 | 10827 | 162 | reverse | |
| ORF22 | 11109 | 11309 | 201 | reverse | |
| ORF23 | 11306 | 11575 | 270 | reverse | |
| ORF24 | 11721 | 11915 | 195 | reverse | |
| ORF25 | 11918 | 12472 | 555 | reverse | |
| ORF26 | 12472 | 12984 | 513 | reverse | |
| ORF27 | 13040 | 13699 | 660 | reverse | |
| ORF28 | 13696 | 14133 | 438 | reverse | |
| ORF29 | 14109 | 14396 | 288 | reverse | |
| ORF30 | 14425 | 14715 | 291 | reverse | |
| ORF31 | 14771 | 15205 | 435 | reverse | |
| ORF32 | 15217 | 15321 | 105 | reverse | |
| ORF33 | 15305 | 15808 | 504 | reverse | |
| ORF34 | 15819 | 16310 | 492 | reverse | |
| ORF35 | 16312 | 16815 | 504 | reverse | |
| ORF36 | 16833 | 17114 | 282 | reverse | |
| ORF37 | 17116 | 17607 | 492 | reverse | |
| ORF38 | 17614 | 17886 | 273 | reverse | |
| ORF39 | 18211 | 18489 | 279 | reverse | |
| ORF40 | 18621 | 18893 | 273 | forward | |
| ORF41 | 18883 | 19143 | 261 | reverse | |
| ORF42 | 19275 | 19544 | 270 | reverse | |
| ORF43 | 19576 | 20238 | 663 | reverse | |
| ORF44 | 20235 | 20450 | 216 | reverse | |
| ORF45 | 20453 | 21103 | 651 | reverse | |
| ORF46 | 21100 | 21627 | 528 | reverse | **Putative SLOG family protein** |
| ORF47 | 21617 | 21943 | 327 | reverse | |
| ORF48 | 21933 | 22127 | 195 | reverse | |
| ORF49 | 22349 | 22540 | 192 | reverse | |
| ORF50 | 22540 | 23028 | 489 | reverse | |
| ORF51 | 23126 | 23482 | 357 | reverse | **Putative CHC2 zinc finger containing protein** |
| ORF52 | 23482 | 24684 | 1203 | reverse | |
| ORF53 | 24728 | 24961 | 234 | reverse | |
| ORF54 | 24951 | 27596 | 2646 | reverse | |
| ORF55 | 28420 | 29502 | 1083 | forward | |
| ORF56 | 29499 | 32120 | 2622 | forward | |
| ORF57 | 32187 | 32435 | 249 | forward | |
| ORF58 | 32445 | 34034 | 1590 | forward | **Putative helicase** |
| ORF59 | 34031 | 34768 | 738 | forward | |
| ORF60 | 34780 | 35904 | 1125 | forward | **Puttive nuclease activity** |
| ORF61 | 35915 | 36361 | 447 | forward | |
| ORF62 | 36416 | 37003 | 588 | forward | **Putative nucleotide pyrophosphorylase** |

| | | | | | |
|---|---|---|---|---|---|
| ORF63 | 36981 | 37802 | 822 | forward | |
| ORF64 | 37807 | 38370 | 564 | forward | **Putative chaperone** |
| ORF65 | 38384 | 40072 | 1689 | forward | **Putative chaperone** |
| ORF66 | 40221 | 40373 | 153 | forward | |
| ORF67 | 40409 | 43393 | 2985 | forward | **Putative DNA polymerase** |
| ORF68 | 43583 | 44200 | 618 | forward | |
| ORF69 | 44233 | 47292 | 3060 | reverse | **Putative tail sheath** |
| ORF70 | 47362 | 47493 | 132 | reverse | |
| ORF71 | 47608 | 50403 | 2796 | reverse | |
| ORF72 | 50535 | 52541 | 2007 | reverse | **Putative helicase** |
| ORF73 | 52704 | 53246 | 543 | reverse | **Putative dihydrofolate reductase** |
| ORF74 | 53331 | 54722 | 1392 | reverse | **Putative helicase/primase** |
| ORF75 | 54739 | 55476 | 738 | reverse | |
| ORF76 | 55593 | 57365 | 1773 | reverse | |
| ORF77 | 57375 | 58094 | 720 | reverse | |
| ORF78 | 58097 | 58258 | 162 | reverse | **Putative chaperone** |
| ORF79 | 58275 | 59636 | 1362 | reverse | **Putative helicase** |
| ORF80 | 59691 | 60998 | 1308 | reverse | |
| ORF81 | 61040 | 61273 | 234 | forward | |
| ORF82 | 61275 | 61955 | 681 | reverse | |
| ORF83 | 62001 | 62615 | 615 | reverse | **Putative baseplate wedge protein** |
| ORF84 | 62619 | 63272 | 654 | reverse | **Putative DNA end protecting protein** |
| ORF85 | 63316 | 63699 | 384 | forward | |
| ORF86 | 63758 | 64360 | 603 | reverse | |
| ORF87 | 64320 | 64523 | 204 | reverse | |
| ORF88 | 64532 | 64810 | 279 | reverse | |
| ORF89 | 64820 | 65260 | 441 | reverse | |
| ORF90 | 65321 | 65791 | 471 | reverse | |
| ORF91 | 65805 | 66467 | 663 | reverse | |
| ORF92 | 66471 | 67625 | 1155 | reverse | **Putative thymidylate synthetase** |
| ORF93 | 67707 | 68411 | 705 | reverse | |
| ORF94 | 68586 | 69206 | 621 | reverse | |
| ORF95 | 69188 | 70024 | 837 | reverse | **Putative guanylate kinase** |
| ORF96 | 70098 | 71411 | 1314 | reverse | |
| ORF97 | 71500 | 72132 | 633 | forward | **Putative hydrolase** |
| ORF98 | 72167 | 72355 | 189 | forward | |
| ORF99 | 72339 | 72542 | 204 | reverse | |
| ORF100 | 72549 | 72845 | 297 | reverse | |
| ORF101 | 72845 | 73624 | 780 | reverse | **Putative baseplate protein** |
| ORF102 | 73699 | 75603 | 1905 | forward | |
| ORF103 | 75645 | 77216 | 1572 | reverse | **Putative baseplate wedge protein** |
| ORF104 | 77246 | 78010 | 765 | reverse | **Putative endonuclease** |
| ORF105 | 78049 | 80061 | 2013 | forward | |
| ORF106 | 80063 | 83206 | 3144 | forward | |
| ORF107 | 83270 | 85345 | 2076 | forward | |
| ORF108 | 85417 | 86064 | 648 | reverse | |
| ORF109 | 86100 | 86387 | 288 | reverse | |
| ORF110 | 86395 | 86685 | 291 | reverse | |
| ORF111 | 86688 | 86939 | 252 | reverse | |
| ORF112 | 86943 | 87311 | 369 | reverse | |
| ORF113 | 87413 | 87964 | 552 | reverse | |
| ORF114 | 87964 | 88302 | 339 | reverse | |
| ORF115 | 88389 | 91067 | 2679 | forward | |
| ORF116 | 91070 | 91561 | 492 | forward | **Putative head completion protein** |
| ORF117 | 91554 | 91925 | 372 | reverse | |
| ORF118 | 91950 | 93131 | 1182 | reverse | **Putative DNA primase/helicase** |
| ORF119 | 93234 | 94436 | 1203 | reverse | |
| ORF120 | 94589 | 94993 | 405 | reverse | |
| ORF121 | 95038 | 95433 | 396 | reverse | |
| ORF122 | 95698 | 96267 | 570 | reverse | |
| ORF123 | 96272 | 99046 | 2775 | reverse | **Putative nudix hydrolase** |
| ORF124 | 99059 | 99814 | 756 | reverse | |
| ORF125 | 99826 | 100794 | 969 | reverse | |

| | | | | | |
|---|---|---|---|---|---|
| ORF126 | 100791 | 101453 | 663 | reverse | |
| ORF127 | 102287 | 102688 | 402 | forward | |
| ORF128 | 102636 | 104255 | 1620 | forward | |
| ORF129 | 104308 | 104769 | 462 | forward | |
| ORF130 | 104766 | 105500 | 735 | forward | |
| ORF131 | 105670 | 105957 | 288 | forward | **Putative DNA binding protein** |
| ORF132 | 106029 | 106430 | 402 | reverse | |
| ORF133 | 106442 | 109003 | 2562 | reverse | |
| ORF134 | 109068 | 109832 | 765 | forward | **Putative neck protein** |
| ORF135 | 109839 | 110087 | 249 | reverse | |
| ORF136 | 110089 | 111072 | 984 | reverse | **Putative sliding clamp protein** |
| ORF137 | 111157 | 111639 | 483 | reverse | **Putative peptidase/plate protein** |
| ORF138 | 111742 | 112755 | 1014 | forward | **Putative ribonucleoside reductase beta** |
| ORF139 | 112888 | 113595 | 708 | forward | |
| ORF140 | 113692 | 117906 | 4215 | forward | **Putative prohead scaffold** |
| ORF141 | 118051 | 119394 | 1344 | forward | **Putative adenylosuccinate synthetase** |
| ORF142 | 119418 | 121289 | 1872 | forward | **Putative terminase large subunit** |
| ORF143 | 121438 | 123144 | 1707 | forward | **Putative ribonucleoside reductase alpha** |
| ORF144 | 123173 | 123388 | 216 | forward | |
| ORF145 | 123471 | 125399 | 1929 | forward | **Putative portal protein** |
| ORF146 | 125545 | 127206 | 1662 | forward | **Putative major capsid protein** |
| ORF147 | 127316 | 127843 | 528 | reverse | **Putative baseplate wedge protein** |
| ORF148 | 127955 | 129646 | 1692 | reverse | |
| ORF149 | 129735 | 131090 | 1356 | forward | **Putative adenylosuccinate lyase** |
| ORF150 | 131087 | 131665 | 579 | reverse | |
| ORF151 | 131717 | 132250 | 534 | forward | |
| ORF152 | 132257 | 133225 | 969 | forward | **Putative endopeptidase** |
| ORF153 | 133276 | 136560 | 3285 | reverse | |
| ORF154 | 136761 | 137570 | 810 | forward | |
| ORF155 | 137610 | 138587 | 978 | forward | |
| ORF156 | 138580 | 138750 | 171 | forward | |
| ORF157 | 138798 | 139484 | 687 | forward | |
| ORF158 | 139481 | 139900 | 420 | reverse | |
| ORF159 | 139939 | 140550 | 612 | reverse | |
| ORF160 | 140555 | 141133 | 579 | reverse | **Putative endonuclease** |
| ORF161 | 141133 | 143094 | 1962 | reverse | **Putative DNA ligase** |
| ORF162 | 143181 | 143567 | 387 | reverse | **Putative endonuclease** |
| ORF163 | 143571 | 144095 | 525 | reverse | **Putative glycoside hydrolase** |
| ORF164 | 144189 | 144629 | 441 | reverse | |
| ORF165 | 144633 | 145253 | 621 | reverse | |
| ORF166 | 145241 | 145546 | 306 | reverse | |
| ORF167 | 145284 | 145433 | 150 | forward | |
| ORF168 | 145461 | 145643 | 183 | forward | |
| ORF169 | 145664 | 146389 | 726 | reverse | |
| ORF170 | 146422 | 147570 | 1149 | reverse | |
| ORF171 | 147563 | 147745 | 183 | reverse | |
| ORF172 | 147757 | 151512 | 3756 | reverse | |
| ORF173 | 151538 | 151645 | 108 | reverse | |
| ORF174 | 151666 | 151965 | 300 | reverse | |
| ORF175 | 151969 | 153396 | 1428 | reverse | **Putative leucine rich repeat protein** |
| ORF176 | 153401 | 155008 | 1608 | reverse | **Putative leucine rich repeat protein** |
| ORF177 | 155023 | 159075 | 4053 | reverse | |
| ORF178 | 159186 | 159737 | 552 | reverse | |
| ORF179 | 159766 | 160479 | 714 | reverse | |
| ORF180 | 160470 | 160751 | 282 | reverse | |
| ORF181 | 160820 | 161338 | 519 | reverse | |
| ORF182 | 161494 | 161748 | 255 | reverse | |
| ORF183 | 161748 | 162197 | 450 | reverse | |
| ORF184 | 162172 | 162672 | 501 | reverse | |
| ORF185 | 162706 | 163071 | 366 | reverse | |
| ORF186 | 163077 | 163442 | 366 | reverse | |
| ORF187 | 163461 | 163820 | 360 | reverse | |
| ORF188 | 163829 | 164506 | 678 | reverse | |

| | | | | |
|---|---|---|---|---|
| ORF189 | 164503 | 165036 | 534 | reverse |
| ORF190 | 165020 | 165349 | 330 | reverse |
| ORF191 | 165367 | 165699 | 333 | reverse |
| ORF192 | 165701 | 166381 | 681 | reverse |
| ORF193 | 166493 | 167233 | 741 | reverse |
| ORF194 | 167243 | 167563 | 321 | reverse |
| ORF195 | 167562 | 167711 | 150 | forward |
