## Supporting information file 1 for "New Finnlakevirus isolate FLiP-2 provides insight into the ecology of ssDNA phages in Flavobacterium hosts"

### PCR details:

**Primer sequences (5'-3'):**

| | |
|---|---|
| FLiP MCP Forward | GAATGTTGTTCGCGGTGCTT |
| FLiP MCP Reverse | CGACCAATGGGAAGAGGGAG |
| FLiP Rep Forward | TCAGCGCAAAGGTTAGGCAT |
| FLiP Rep Reverse | GCTGTGCTAACGCCCAAATC |
| FLiP genome Forward | GCGCAAAGGTTAGGCATAGC |
| FLiP genome Reverse | AACTTTACCGCTATCGCCGT |

**PCR program (MCP and Rep):**

| | | |
|---|---|---|
| 95°C | 7 min | |
| 95°C | 30 s | 30x |
| 62°C | 30 s | 30x |
| 72°C | 1 min | 30x |
| 72°C | 10 min | |
| 12°C | ∞ | |

**PCR Program (genome):**

| | | |
|---|---|---|
| 98°C | 30 s | |
| 98°C | 10 s | 30x |
| 64.8°C | 30 s | 30x |
| 72°C | 4 min. 36 s. | 30x |
| 72°C | 10 min. | |
| 12°C | ∞ | |

Table S1. Water samples collected from freshwaters in Finland and filtered through 0.2 µm filters. PCR analysis of the DNA extracted from filter membranes using primers targeting either FLiP major capsid protein (MCP) or replication initiation protein (Rep) genes. All positive results are included, even when the PCR product size diverges from respective product size of FLiP.

| Name of lake or river | Municipality | Date | Positive in PCR | |
|---|---|---|---|---|
| | | | **MCP** | **Rep** |
| Vuojärvi | Laukaa | 26.8.2019 | + | - |
| Vuojärvi | Laukaa | 22.8.2022 | + | - |
| Saraavesi | Laukaa | 26.8.2019 | - | - |
| Saraavesi | Laukaa | 22.8.2022 | + | - |
| Peurunkajärvi | Laukaa | 26.8.2019 | + | + |
| Peurunkajärvi | Laukaa | 22.8.2022 | + | - |
| Vuonteensalmi | Laukaa | 18.9.2019 | + | + |
| Vuonteensalmi | Laukaa | 22.8.2022 | + | + |
| Kiesimenjärvi | Saarijärvi | 11.8.2019 | - | - |
| Kiesimenjärvi | Saarijärvi | 20.8.2022 | + | - |
| Ilveslahti, Likosalmi | Laukaa | 22.8.2022 | - | - |
| Haapaniemi, Päijänne | Jyväskylä | 22.8.2022 | - | - |
| Tikka, Päijänne | Jyväskylä | 20.8.2022 | + | + |
| Äijälänsalmi | Jyväskylä | 20.8.2022 | + | - |
| Suuruspää, Jyväsjärvi | Jyväskylä | 20.8.2022 | + | - |
| Luonetjärvi | Jyväskylä | 21.8.2022 | + | - |
| Kaivovesi | Jyväskylä | 22.8.2022 | - | - |
| Tuomiojärvi | Jyväskylä | 20.8.2022 | + | - |
| Kuhnamo | Äänekoski | 22.8.2022 | - | - |
| Äänejärvi | Äänekoski | 22.8.2022 | + | - |
| Mansikkaniemi | Saarijärvi | 20.8.2022 | - | - |
| Pielinen | Lieksa | 2.8.2019 | + | - |
| Tielampi | Lapinlahti | 3.8.2019 | + | - |
| Oulujärvi | Vaala | 4.8.2019 | + | + |
| Särkinen | Vaala | 4.8.2019 | + | - |
| Pantiolampi | Vaala | 4.8.2019 | - | - |
| Haarusjärvi | Kauhava | 5.8.2019 | + | + |
| Iiruunjärvi | Alajärvi | 6.8.2019 | + | - |
| Kankarisvesi | Jämsä | 8.8.2019 | + | - |
| Jämsänjoki | Jämsä | 8.8.2019 | + | - |
| Ahvenlampi | Laukaa | 16.8.2019 | + | - |
| Elänne | Mänttä-Vilppula | 25.8.2019 | - | - |

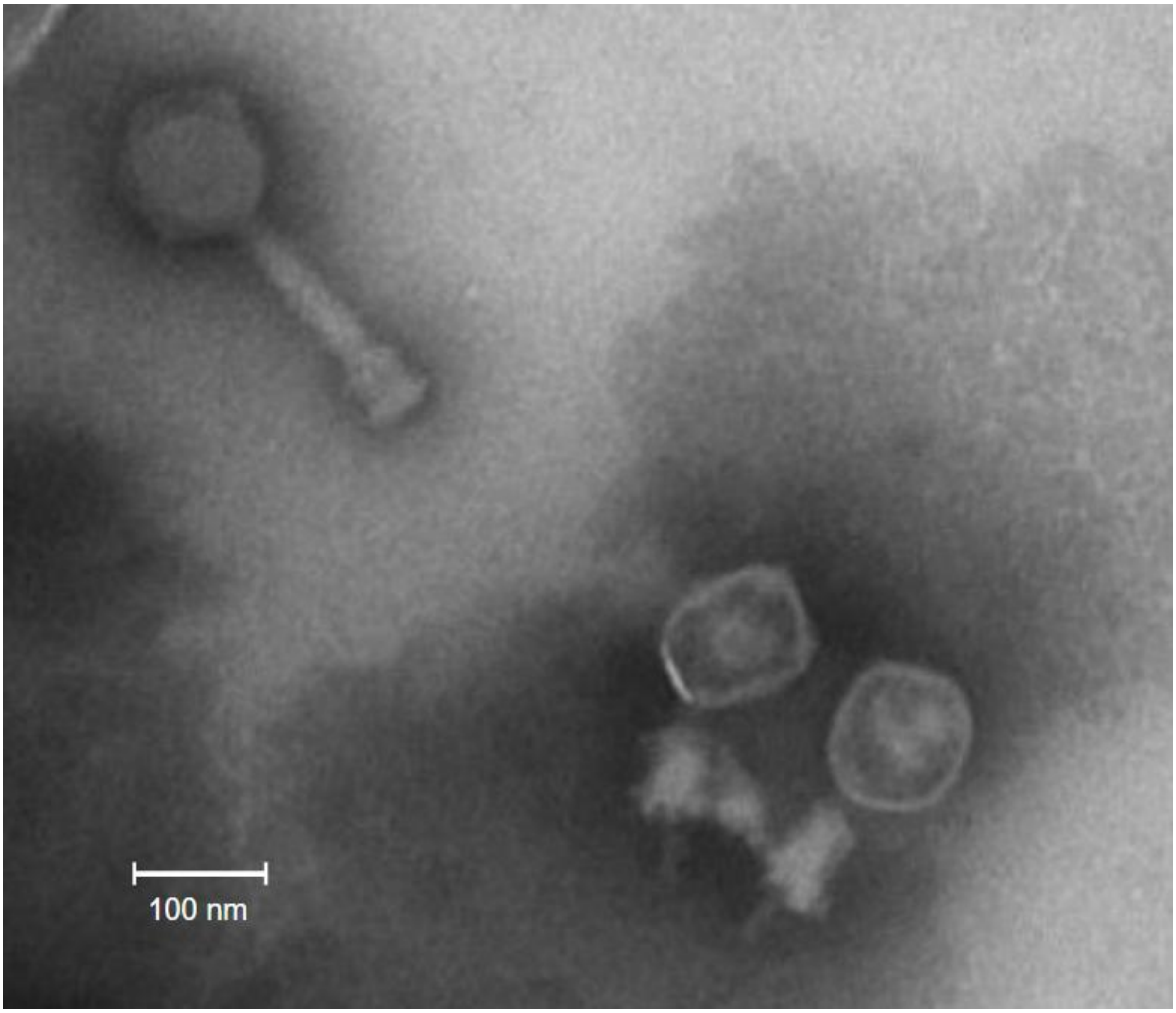


Figure S1. EM image of MaF61 particles. In the upper left corner, a phage particle with an uncontracted tail is visible, while in the lower right, two particles with contracted tails can be observed. Scale bar is 100 nm.

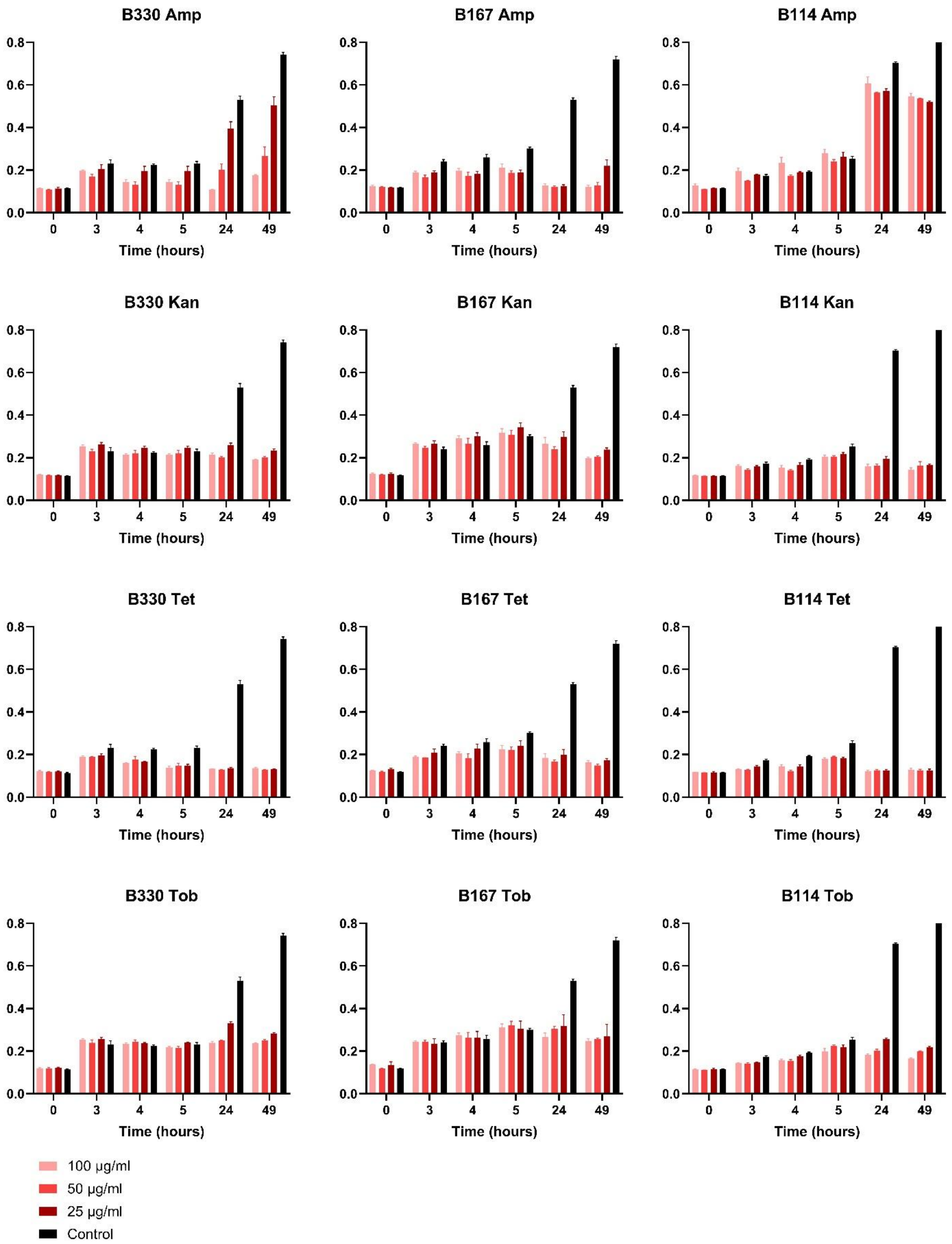


Figure S2. Effect of ampicillin (Amp), kanamycin (Kan), tetracycline (Tet) and tobramycin (Tob) to growth of Flavobacterium sp. B330, B167 and B114.
